# Sleep loss impairs myelin function by altering cholesterol metabolism in oligodendroglia

**DOI:** 10.1101/2023.11.27.568716

**Authors:** Reyila Simayi, Eleonora Ficiarà, Oluwatomisin Faniyan, Antonio Cerdán Cerdá, Amina Aboufares El Alaoui, Rosamaria Fiorini, Adele Cutignano, Fabiana Piscitelli, Pablo Andújar, Alexandra Santos, Federico Del Gallo, Luisa de Vivo, Silvia De Santis, Michele Bellesi

## Abstract

The increasing prevalence of sleep deprivation in modern society demands comprehension of the biological consequences of sleep loss on brain functioning. Our study reveals significant effects of sleep deprivation on myelin integrity. As a result, we identify increased conduction delays in nerve signal propagation, hindered interhemispheric synchronization, and impaired motor performance associated with sleep loss. By profiling oligodendrocyte transcriptome and lipidome, we observe sleep loss-induced endoplasmic reticulum stress and lipid metabolism disruption, particularly affecting cholesterol homeostasis. This shift in cholesterol levels alters myelin physical properties. Boosting cholesterol transport to myelin sheaths prevents sleep loss effects on nerve signal propagation and behavior. Our findings highlight the critical role of oligodendrocyte cholesterol regulation in behavioral deficits associated with sleep loss and unveil a novel target for intervention.

## INTRODUCTION

While the functions of sleep remain to be fully understood, it is clear that even a single missed night of sleep or consistently shortened sleep durations over extended periods can importantly affect brain functioning and behavior. Slow reaction times and increased errors of omission at the psychomotor vigilance task (PVT) are the most objective changes in alertness associated with sleep loss (*1*, *2*). These changes usually coincide with sleep episodes intruding into wakefulness and a slowdown of waking electroencephalographic (EEG) rhythms (*3*).

However, how sleep loss causes behavioral impairment is unclear. Considerable work has been undertaken to determine the characteristics of the ‘neural fatigue’ and to uncover the mechanisms causing neural network anomalies that result in compromised behavior. For instance, it has been found that sleep loss can induce brief periods of neuronal silence (i.e., local off periods) in an otherwise awake rat brain. This cellular effect has been related to impaired motor performance in a food-pellet reaching task during sleep deprivation (*4*). Similarly, in humans who had intracranial electrodes implanted to monitor individual neurons, there were noticeable local changes in neuronal activity just before attention lapses occurred during the PVT (*5*).

While most of the attention has been focused on neurons, other brain cells have received little consideration, despite the evidence showing that glial cells can respond to sleep loss with molecular changes and extensive structural modifications (*6*, *7*). For instance, electron microscopy investigations in mice have shown that chronic sleep restriction can lead to decreased myelin thickness and changes in the nodal region’s geometry of callosal axons, implicating adaptive remodeling of oligodendrocytes (*8*, *9*). Here, we explore the effects of sleep loss on oligodendrocyte function, investigating its impact on neuronal signal conduction, synchronization, and its contribution to the behavioral deficits associated with sleep deprivation.

## RESULTS

### Sleep loss induces widespread white matter changes in the rat and human brain

Poor sleep quality and short sleep duration have been associated with magnetic resonance imaging (MRI) alterations in white matter (WM) microstructure in multiple brain regions (*10–12*). In the first analysis, we sought to confirm this association in a large sample of healthy individuals (n=185) by quantifying the relation between WM microstructural integrity and the Pittsburgh Quality Index (PSQI), a well-recognized metric of sleep quality. We found a significant negative correlation between the PSQI and MRI makers of WM integrity indicating that poor sleep quality was related to lower microstructural integrity. Such effect was more pronounced and widespread in the WM skeleton when microstructural integrity was assessed using the fractional anisotropy (FA, Fig. 1A). Other structural markers such as the restricted function (RF) derived from the CHARMED model (*13*) also showed a significant but weaker association with PSQI (Supplementary results, fig. S1, S2).

**Fig. 1:**
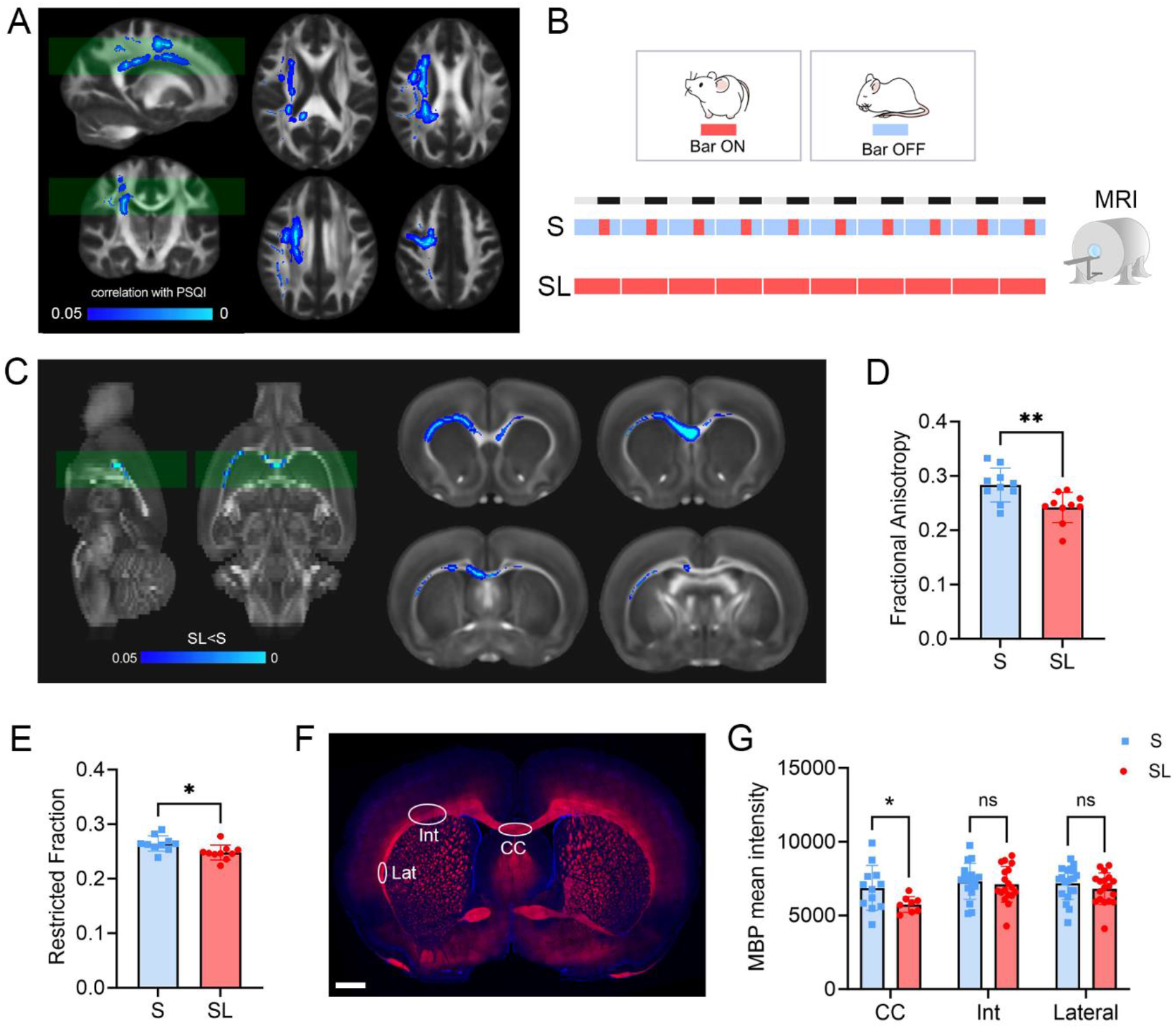
Widespread and inter-species effects of SL on brain WM. **A**. Significant association between fractional anisotropy (FA) and PSQI in the HCP database (n=185), superimposed on the FA template used in the analysis. Blue-light blue: significant negative association (p<0.05). The opposite contrast is not significant. **B**. Rat experimental design (sleep [S], n=10; sleep loss [SL], n=10). **C.** Significant differences of FA in SL relative to S, superimposed on the FA template used in the analysis. Blue-light blue: significant reduction in SL (p<0.05). The opposite contrast is not significant. **D-E**. Quantification of FA (**D**) and RF (**E**) in S (blue) and SL (red) rats. *P<0.05; **P<0.01. **F**. Example of MBP staining (red) and DAPI (blue) in a S rat frontal section. White circles indicate where MBP staining was quantified. Scale Bar: 2mm **G**. Quantification of MBP staining in corpus callosum (CC), intermediate (Int), and lateral (Lateral) subcortical WM. *P<0.05. ns: not significant.

To confirm the causal link between WM microstructural integrity and sleep disruption, in a second experiment we measured WM microstructural integrity in rats that had been subjected to sleep restriction (sleep loss [SL] group) and in matched normal sleeping controls (S) using structural MRI. We first validated with polysomnography an automatic sleep restriction approach (see supplementary results) and then we used it to sleep restrict rats for 10 days prior the MRI scanning (Fig. 1B). Voxel wise analysis of the WM revealed a widespread decrease in FA and RF in SL animals relative to controls, which was indicative of reduced microstructural integrity due to SL (Fig. 1C-E).

MRI structural markers do not possess the specificity to distinguish whether the loss of structural integrity is due to myelin or axonal impairments. However, previous ultrastructural analysis of more than 17000 axons of mouse corpus callosum and lateral olfactory tract has demonstrated that SL can specifically reduce myelin thickness without a significant change in axonal density and axon diameter (*8*). To confirm the effect of SL on myelin in this dataset, after the MRI scan brains were stained with myelin basic protein (MBP), a protein highly expressed in myelin sheaths (Fig. 1F). Microscopic quantification of MBP immunoreactivity in the same brain regions reporting a FA decrease revealed a significant reduction of MBP immunoreactivity in SL rats relative to S in the corpus callosum (P=0.045). The effect of SL on MBP expression in lateral portions of subcortical WM was not significant (P=0.84 for Intermediate, P=0.55 for Lateral, Fig. 1G).

### Sleep loss increases transcallosal evoked LFP latencies

While both ultrastructural and MRI findings hint at changes in myelin structure after sleep deprivation, whether myelin function is also altered in vivo has never been tested. Based on MRI and ultrastructural studies, we postulated that sleep loss-induced myelin modifications could result in delayed signal transmission, i.e., reduced speed of action potentials travelling along the axons. Therefore, we measured in vivo the latency of transcallosal cortical local field potentials (LFPs) evoked by the electrical stimulation of the contralateral homotopic cortex, which is inversely related to the signal conduction velocity along the connections. To evaluate changes in response latency as a function of sleep and SL, we implanted two groups of rats with bipolar concentric electrodes for electrical stimulation and chronic intracortical LFP recordings (Fig. 2A). We recorded LFPs from the left frontal cortex after electrical stimulation of the right frontal cortex before (baseline) and after sleep manipulation (post session, Fig. 2B). We then measured the latency of the first negative component of the transcallosal evoked response (Fig. 2B). By comparing the post session with the baseline, we found that the latency of the early negative component was delayed by 32.8±27.3% (P<0.0001) in the SL group, while it did not change in S rats (P=0.99), thus confirming SL-dependent increase in conduction delay in vivo (Fig. 2C-D). Increased conduction delays could also modify the amplitude and the slope of the evoked potential, especially if the decrease in myelination is heterogeneous among callosal bundles. However, despite the large variability among the animals, neither the amplitude nor the slope of the evoked response significantly changed after SL (Amplitude: P=0.81, Slope: P=0.96, Fig. S4A-D).

**Fig. 2.**
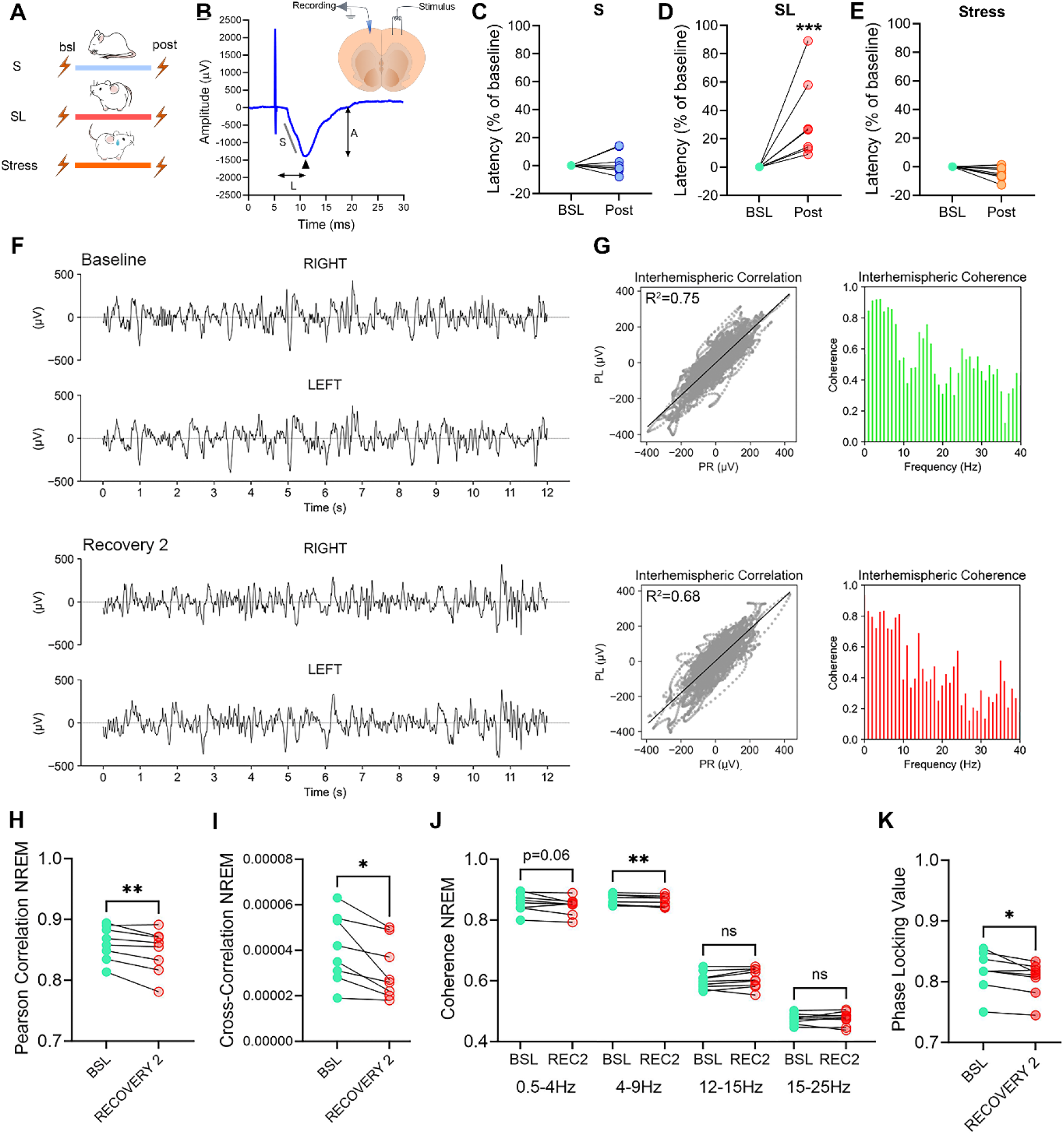
SL increases conduction delays and reduces interhemispheric synchronization. **A**. Experimental design. **B**. Example of an evoked cortico-cortical transcallosal response in a S rat. L: peak latency; A: peak amplitude; S: slope. Arrows indcates the negative peak of the early component of the evoked response. The picture above indicates the location of the stimulation and recording LFP electrodes in the frontal cortex. **C-E**. Mean latency values of the early component negative peak for S (**C**, blue, n=8), SL (**D**, red, n=8), and Stress (**E**, orange, n=8) individual rats. Values are represented as % relative to their own baseline. ***P<0.001. **F.** two examples of raw 12-s EEG records of NREM sleep from the right and left parietal derivations for a SL rat during baseline (upper panel) and during the second day of recovery after sleep restriction (lower panel). **G**. Left panels: voltage values of the left derivation were plotted as a function of the right derivation for corresponding consecutive sampling intervals for the 12-s records depicted in **F**. The bold line represents a linear regression. Right panels: interhemispheric coherence spectra for frequencies between 0.5 and 40 Hz of the corresponding 12-s records depicted in **F**. **H-I**. Interhemispheric correlation (**H**) and cross-correlation (**I**) values for SL rats (n=6) computed for baseline and for the second day of recovery after sleep restriction (recovery 2). *P<0.05. **P<0.01. **J**. Interhemispheric coherence values for delta (0.5-4 Hz), theta (4-9 Hz), sigma (12-15 Hz), and beta (15-25 Hz) power bands computed for baseline and for recovery 2 (REC2). **P<0.01. **K**. Interhemispheric phase-locking values for SL rats computed for baseline and for recovery 2.

While we intentionally used automated sleep restriction to reduce hands-on interaction with the animals, sleep deprivation is broadly recognized as a stressful condition, often elevating stress markers like corticosterone levels (*8*). Additionally, there is evidence suggesting that stress can negatively affect myelin (*14*). To understand whether the observed functional effects on myelin were related to the stress from the sleep restriction procedure, we evaluated cortico-cortical evoked responses in a separate rat group before and after subjecting them to 10 days of unpredictable stress. This approach, involving a mix of varied and random stressors, is a standard method to study stress effects. The analysis of the evoked responses did not reveal any difference in latency, amplitude, or slope between baseline and post stress sessions (Latency: P=0.56; Amplitude: P=0.5; Slope: P=0.99; Fig. 2E, Fig. S4E-F), effectively ruling out stress as a confounding factor in the observed increased conduction latencies post-SL.

### Sleep loss impairs interhemispheric synchronization of neuronal activity and motor behavior

We have shown that SL can result in a reduced nerve pulse propagation speed compared to normal sleeping animals. Therefore, we posited that this could cause broad delays in the timing of action potentials (*15*), especially for those travelling on the heavily myelinated fibers of the corpus callosum. As a result, motor coordination that relies on synchronous interhemispheric activity could be altered by the increased callosal conduction delay. To test this hypothesis, we first verified whether interhemispheric synchronization of neuronal activity was affected by SL. We measured the correlation and coherence of the EEG signals between the two hemispheres during 24h of baseline and after 4 days of sleep restriction (Fig. 2F-G, S3A). We decided to focus this analysis primarily in NREM sleep, which is characterized by a low arousal tone. We reasoned that the activation of the arousal systems typically observed in activated states such as wake and REM sleep, could boost interhemispheric coherence given their bi-hemispheric bottom-up projections. Indeed, we found that the temporal correlation was reduced during NREM sleep, while it was unchanged during wake and REM sleep (NREM: Pearson’s correlation: P=0.0095; cross-correlation: P=0.011. Wake: Pearson’s correlation: P=0.76; cross-correlation: P=0.36. REM: Pearson’s correlation: P=0.41; cross-correlation: P=0.35, Fig. 2H-I, Fig. S3A-B). Further analysis revealed a general decrease of coherence in the frequency domain of delta and theta (P=0.06 for delta, P=0.01 for theta) and a lack of effect for the other faster frequencies (P=0.22 for sigma; P=0.95 for beta, Fig. 2J). These effects were associated with a reduction of the phase locking value computed for delta-theta frequencies (P=0.02, Fig. 2K).

We then exposed another set of rats to 10 days of sleep restriction as in previous experiments, and subsequently assessed their motor skills on a rotarod apparatus. Relative to normal sleeping rats, sleep restricted rats showed diminished motor performance whilst motor learning was largely unaffected (motor performance: P=0.0037, motor learning: P=0.21, Fig. 5D-F). Thus, SL impaired interhemispheric synchronization of neuronal activity and led to motor performance deficits.

### Sleep loss induces ER stress and affects lipid metabolism in oligodendrocytes

Morphological and functional data suggest that SL can lead to myelin deficits, which may be attributed to an overall dysfunction of the oligodendroglia. For a more in-depth understanding, we interrogated a publicly available gene dataset that was previously obtained in 2’,3’-cyclicnucleotide3’-phosphodiesterase(CNP)-eGFP-L10a mice using TRAP technology combined with microarray analysis to tag polysomes and immunoaffinity purify oligodendrocyte specific mRNAs (*6*). By comparing data from sleeping mice (S, 6–7 h of sleep during the light phase; n = 6) with data obtained from sleep deprived mice (4 h of SL during the light phase through exposure to novel objects, n = 6, Fig. 3A), we identified 5035 probe sets differentially expressed because of SL (8.9% of 45101 probe sets; false discovery rate = 1%), representing 3448 unique genes, including 1889 upregulated and 1559 downregulated genes (Fig. 3B). Upregulated and downregulated genes were clustered to distinct biological processes using the gene annotation enrichment analysis (Fig. 3C). Among the upregulated genes, an importantly enriched class was related to endoplasmic reticulum (ER) stress. Specifically, we found genes associated with PERK and IRE1 arms of the unfolding protein response (e.g., Xbp1, Hif3a, Eif2ak4) and to the ER associated degradation (ERAD) such as Hspa1a, Hsp90aa1, and Hsph1. Other upregulated genes were involved in lipid degradation of plasma membrane phospholipids (e.g., Plcb1 and Plcd4) and lysophospolipids (e.g., Naaa, Abhd4, Asp, Lipa). The most enriched classes of downregulated genes were related to lipid biosynthesis and trafficking, which included genes involved in the synthesis of glycerophospholipids (e.g., Digat2, Lpcat2, Elovl7) and cholesterol (e.g., Dhcr7, Msmo1, Mvk). In summary, the analysis of gene expression in oligodendrocytes revealed that SL significantly affected ER and lipid homeostasis.

**Fig. 3.**
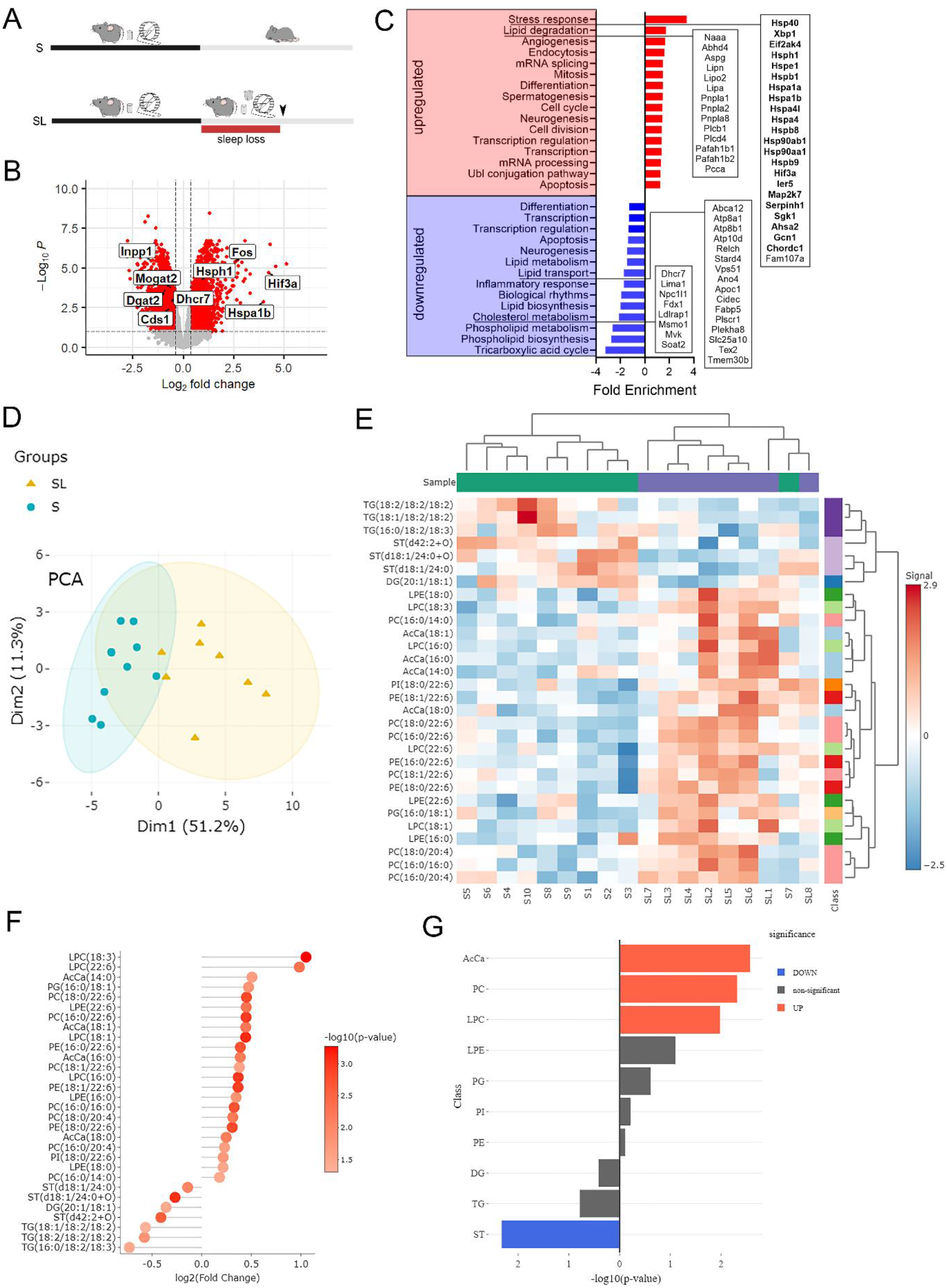
SL induces transcriptional changes in oligodendrocytes and modifies the lipid composition of myelin. **A.** Experimental design of transcriptomics and lipidomic studies. **B**. Vulcan plot of the significant up-regulated and down-regulated transcripts (red) because of SL (S, n=6; SL, n=6). Some transcripts related to ER stress and lipid homeostasis are highlighted in bold. **C**. Functional enrichment analysis identifying significant and highly enriched categories of up-regulated (red) and down-regulated (blue) genes. Categories of up (red) and down-regulated (blue) genes are ranked according to their enrichment score. Genes belonging to some key categories are highlighted in the frames. Note that in the stress response category genes related to ER stress are in bold. **D.** Principal component analysis (PCA) correctly separates the two experimental groups based on lipid expression (S, n=10; SL, n=8). **E**. Hierarchical clustering showing the separation of the S and SL samples according to their differentially expressed lipids. **F**. Distribution of significant up-regulated (right) and down-regulated (left) lipids. Lipids are ranked according to their fold-change, the scale indicates their significance. Acylcarnitine (AcCa), diacylglycerol (DG), lysophosphatidylcholine (LPC), lysophosphatidylethanolamine (LPE), phosphatidylcholine (PC), phosphatidylethanolamine (PE), phosphatidylinositol (PI), sulfatides (ST), triglycerides (TG). **G**. Enrichment analysis identifying the classes of lipids that significantly change because of SL. In red the classes of the up-regulated lipids, while in blue the class of the down-regulated lipids.

### Sleep loss unbalances lipid homeostasis in myelin enriched fractions

Guided by the results of the gene enrichment analysis, we used untargeted lipidomics to characterize the lipid composition of pure myelin fractions derived from mice subjected to 6 hours of SL, through the exposure of novel object during the light cycle, and mice allowed to sleep ad libitum during the same time of the day (S, Fig. 3A). From each mouse forebrain, we derived highly purified myelin fractions that were subjected to lipidomic LC-MS/MS analysis. Principal component analysis (PCA) on the lipidomic data effectively differentiated between the two experimental groups (Fig. 3D). Unsupervised hierarchical clustering unveiled notable distinctions in lipid class composition, particularly an increase in lysophospholipids and glycerophospholipids, as well as a reduction in sulfatides in the SL group (Fig. 3E). To delve deeper into the findings, we conducted a comprehensive investigation of significant lipid species using differential expression analysis (Fig. 3F). An enrichment using over-representation analysis revealed that the significantly upregulated lipids were enriched in the acylcarnitine (AcCa), lysophosphatidylcholine (LPC), and phosphatidylcholine (PC) categories, whereas the down-regulated lipid species exhibited enrichment in the sulfatides (ST) category (Fig. 3G). Thus, SL altered lipid composition of myelin membranes.

### Sleep loss reduces cholesterol levels and alters membrane fluidity in myelin enriched fractions

Given the important downregulation of transcripts related to cholesterol metabolism and the modifications of lipid profiles in myelin enriched fractions, we performed a more narrowly targeted analysis focusing on only brain-specific cholesterol-related processes. We reasoned that this could uncover potential molecular mechanisms regulating cholesterol homeostasis in oligodendrocytes. To highlight cholesterol-related genes that were downregulated, we filtered the GO biological processes annotation terms using the term ‘cholesterol’ and we repeated a differentially expression analysis only in this selective dataset using an FDR of 5%. As illustrated in Fig. 4A, we found that the most notable differences belonged to the GO terms related to cholesterol homeostasis and transport, while biosynthesis and storage were less represented, suggesting a potential impairment of cholesterol intracellular transport and, to a lesser extent, cholesterol production.

**Fig. 4.**
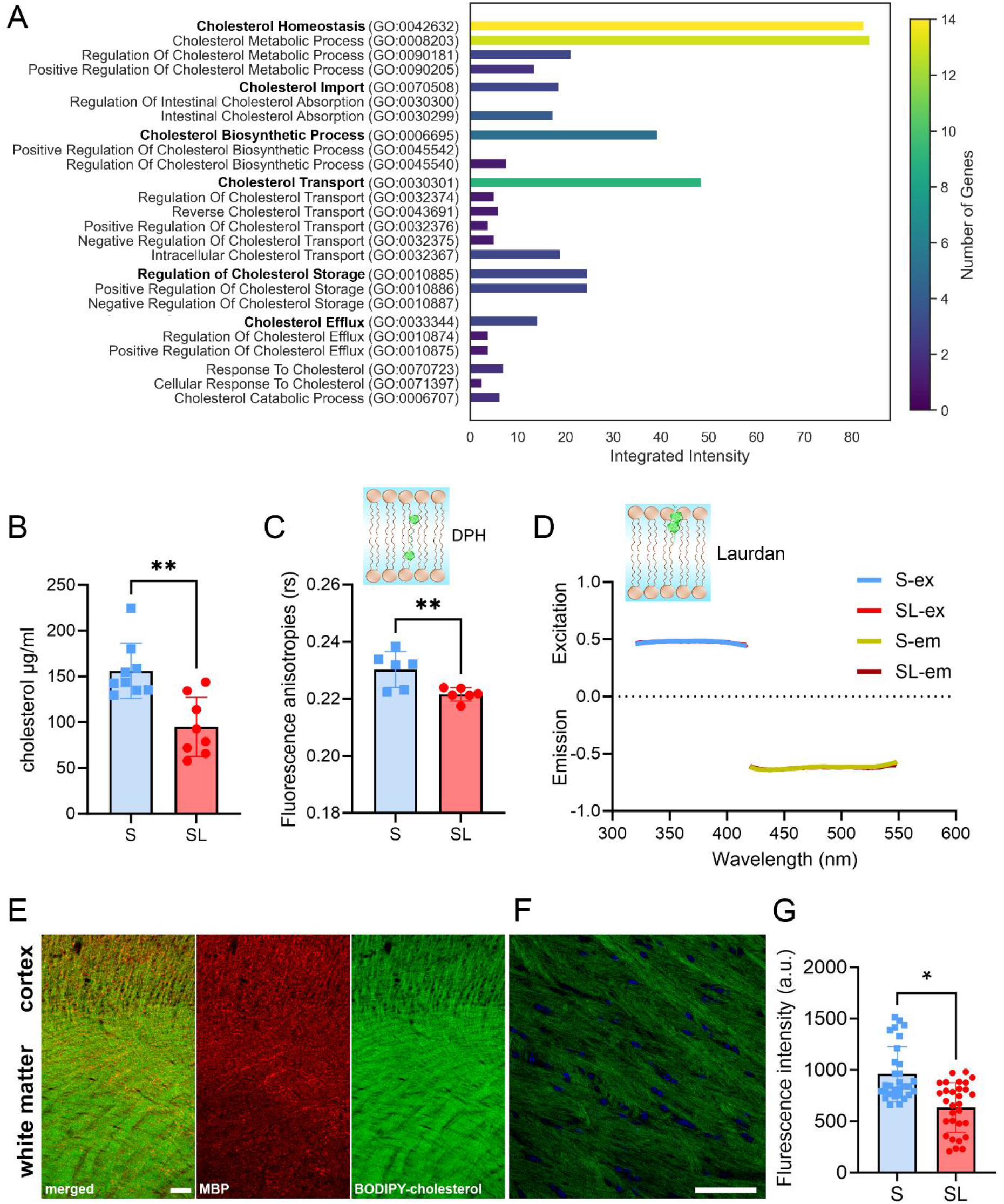
SL alters cholesterol homeostasis affecting myelin membranes fluidity. **A.** Cholesterol-related functional annotation analysis for down-regulated genes. In bold are the ‘parent’ GO terms. Integrated intensity was computed as the sum of the average expression of the genes involved in that specific category. **B.** Cholesterol levels as measured in the purified myelin fractions of S (n=10) and SL (n=8) animals with LC-MS. **P<0.01. **C.** Fluorescence anisotropies inversely related to core membrane fluidity in the purified myelin fractions of S (n=6) and SL (n=6) animals. **P<0.01. **D.** Laurdan excitation and emission spectra measurements in the purified myelin fractions of S and SL animals indicating fluidity stability in the outer part of the myelin membranes. For **C** and **D**, the small inset indicates the location in the plasma membrane where the fluorescent probe binds. **E**. Representative images of BODIPY–cholesterol (green) and anti-myelin basic protein (MBP, red) staining in a rat. Scale bar: 50µm. **F**. Representative image of BODIPY– cholesterol (green) and DAPI (blue) captured in the corpus callosum. Scale bar: 50µm. **G**. Quantification of BODIPY-cholesterol fluorescence intensity in the corpus callosum of S and SL rats. *P<0.05.

**Fig. 5.**
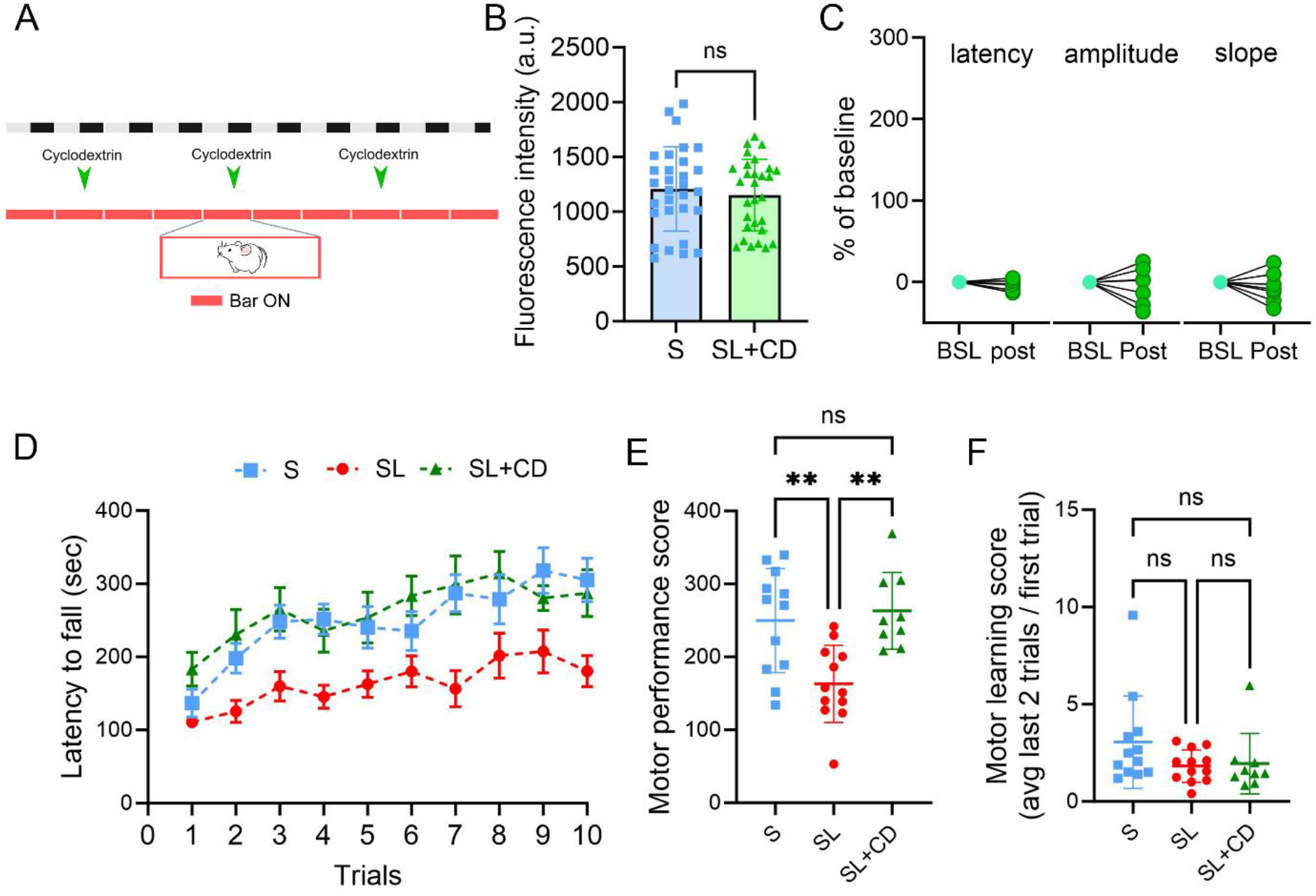
SL-related conduction delays and behavior modifications are prevented by aiding cholesterol transport. **A.** Experimental design. **B**. BODIPY-cholesterol levels in the corpus callosum of S (n=5) and SL+CD (n=5) treated rats. **C.** Mean latency, amplitude, slope values of the early component negative peak for SL individual rats that were treated with Cyclodextrin (n=7). Values are represented as % relative to their own baseline. **D**. Latency to fall (in seconds) during 10 trials for S (n=12), SL (n=12), and SL+CD (n=9) rats of the accelerating rotarod experiment. **E-F**. Motor performance (**E**) and learning (**F**) score of S, SL, and SL+CD rats. **P<0.01.

To determine whether these transcriptional changes correlated with changes in oligodendrocytes cholesterol levels, we conducted LC-MS quantifications on the same myelin enriched fractions previously used for the lipidomic analysis. Quantitative analysis confirmed a significant reduction of cholesterol concentration in SL relative to S mice (P=0.0011, Fig. 4B). Optimal membrane fluidity and curvature in myelin necessitate high cholesterol levels. This ensures membrane stability, minimizes ion leakage, and reinforces the insulating properties of myelin membranes (*16*). To ascertain if the drop in cholesterol correlated with changes in myelin membrane fluidity, we performed spectrofluorimetric assessments using 1,6-diphenyl-1,3,5-hexatriene (DPH), integrated into the hydrophobic lipid section, and 2-dimethylamino-(lauroyl)-naphtalene (Laurdan), positioned at the juncture between the hydrophobic and hydrophilic areas of the membrane. Examination of DPH’s steady-state fluorescence anisotropies (rs) showed a notable rs reduction in myelin derived from the brains of SL mice, pointing to enhanced fluidity in the inner segment of the bilayer (P=0.0097, Figure 4C). However, Laurdan’s quantitative analysis did not exhibit any distinctions between S and SL in both excitation and emission GP spectra (Excitation: P=0.98; Emission: P=0.71, Figure 4D), implying that the fluidity of the external segment of the membrane was not affected by the cholesterol reduction. In summary, these findings indicated that sleep deprivation can increase myelin membrane fluidity by reducing cholesterol levels.

### Sleep loss effects on transcallosal evoked LFP latencies and motor behavior are prevented by aiding cholesterol transport

Both transcriptomic and lipidomic experiments in mice have found that cholesterol homeostasis and transport are the main cellular pathways altered in oligodendrocytes because of SL. We reasoned that by boosting cholesterol transport during SL we could minimize or prevent myelin dysfunctions and restore optimal conduction velocity in rats. Thus, we first confirmed that sleep restriction could reduce cholesterol levels in rats by staining brain tissue from SL and S animals with BODIPY–cholesterol. We focused our analysis on myelin enriched fibers of the corpus callosum, and we found lower levels of fluorescence in SL relative to S, which were indicative of reduced tissue cholesterol concentration (P=0.013, Fig. 4E-G). By contrast, BODIPY–cholesterol levels in astrocytes and neurons were not affected by SL (Fig. S5). Then, we administered 2-hydroxypropyl-β-cyclodextrin (cyclodextrin) to a group of rats and measured its effects on BODIPY–cholesterol levels and on in vivo conduction delay before and after SL. Cyclodextrin is a molecule known to aid cholesterol transport and improve myelination in oligodendroglia (*17*). Three subcutaneous injections of cyclodextrin over 10 days of sleep restriction were able to prevent SL induced reductions in cholesterol levels (P=0.74, Fig. 5A-B) and increases in conduction delay (P=0.83), with no effect on amplitude and slope of the evoked responses (Amplitude: P=0.9, Slope: P=0.91, Fig. 5C). Additionally, cyclodextrin averted the deterioration of motor behavior. Specifically, the performance score in cyclodextrin-treated rats did not change compared to the normal sleeping group (performance P=0.8719, learning P=0.33) but was significantly different from that of SL untreated rats (performance P=0.002, learning P=0.99, Fig. 5D-F), thus suggesting that aiding cholesterol transport and restoring normal myelin functions prevented motor deficits.

## DISCUSSION

We observed a reduction of WM integrity in individuals with poor sleep quality and in rats subjected to sleep restriction. These results validate and broaden earlier observations of myelin ultrastructural changes documented in sleep restricted mice (*8*). The vulnerability of oligodendrocytes to SL may be attributed to the association between sleep deprivation and increased expression of transcripts related to endoplasmic reticulum (ER) stress within oligodendrocytes. Sleep deprivation might intensify the demand for oligodendrocyte resources, amplify metabolism, thereby producing significant oxidative stress, or even result in ER calcium depletion (*18*). It is plausible that ER stress acts as a catalyst, leading to lipid imbalance and subsequent myelin dysfunction. Previous studies have highlighted a link between ER stress and shifts in lipid balance in both oligodendroglia and other cell types (*19–21*), suggesting that lipid homeostasis in oligodendrocytes is finely tuned and may be susceptible to ER stress (*22*).

We found that the transcripts related to the production of proteins crucial for cholesterol synthesis, transport, and overall cholesterol levels, were reduced in myelin enriched fractions. The demonstration that the low cholesterol levels were associated with increased membrane fluidity indicates that the extent of cholesterol reduction induced by SL had a functional impact on the physical property of myelin. Maintaining high cholesterol in myelin membranes is pivotal for preserving natural membrane curvature and optimal insulating capacity (*16*, *23*). By altering the fluidity of myelin membranes, SL could disrupt the insulating function of the myelin membrane and alter the efficiency of saltatory conduction, potentially slowing down nerve impulse transmission.

Indeed, we observed an approximately 33% average increase in the delay of interhemispheric conduction due to SL, although there was considerable variability among individual animals. This phenomenon was associated with decreased synchronization in the activity of homotopic brain regions. Although we were able to confirm this effect specifically during NREM sleep, it is plausible that more nuanced alterations during wakefulness were not detectable through our broad EEG analysis. Computational models suggest that even slight alterations in conduction delay can considerably disrupt network activity, leading to shifts in phase-locked interactions among oscillating networks (*24*). Desynchronized neuronal networks might prompt communication breakdowns between different brain areas, thereby affecting behavior (*25*, *26*). In line with this, we observed decreased motor performance after SL, which may be attributed to changes in myelin and the resulting disruption in network synchronization (*15*).

Importantly, modifications of both signal propagation and behavior were prevented when the animals were treated with cyclodextrin, a drug that promotes cholesterol re-localization to the myelin membranes. These findings establish a causal link between SL, cholesterol homeostasis, signal conduction delays, and behavioral performance, and offers a novel biological mechanism underlying the behavioral impairment associated with the loss of sleep.

## Acknowledgments

We thank Aroa Sanz Maroto for excellent technical support.

## Author contributions

Conceptualization: MB, LdV, SDS. Investigation: RS, EF, OF, AC, AA, RF, ACC, FP, PA, AS, FDG, LdV, SDS, and MB. Writing—original draft: MB, LdV, SDS. Writing—review and editing: All authors. Project supervision and funding: MB, LdV, SDS.

## Data and materials availability

### Data availability

Human MRI data was derived from the latest pre-processing release (v3.19.0) of the Human Connectome Project (HCP database www.humanconnectome.org). Oligodendrocyte transcriptomic data was obtained from NCBI GEO GSE48369 database. Links to remaining raw data will be made available upon publication.

### Code availability

Codes, accompanied by comprehensive guidelines for replicating the analyses discussed in this paper, can be found on GitHub at https://github.com/BSRLab.

## SUPPLEMENTARY MATERIALS

### Materials and Methods

#### Animals

C57BL/6J male mice (postnatal day [P] 40-60) and Wistar rats (P40-80) were used in this study with the exception of gene expression analysis, in which we have referred to a previous database NCBI GEO GSE48369, obtained in adult (9-10 weeks old) heterozygous 2’,3’-cyclicnucleotide 3’-phosphodiesterase (CNP)-eGFP-L10a bacterial artificial chromosome (BAC) transgenic mice of either sex (*6*). All animals were under a light/dark 12:12 with light on at 8 am at a 23 ± 1 °C (environmental temperature) and were provided with food and water available ad libitum and replaced daily at 8 am. All procedures involving animals adhered to the local Institutional Animal Care and Use Committee and the European Communities Council Directives (2010/63/EU, 542/2023-PR) and to the Animals (Scientific Procedures, P5E96A446) Act 1986 and Amendment Regulations 2012 as outlined in UK law and approved by the University of Bristol Animal Welfare and Ethics Review Board.

#### Sleep deprivation

In this study, sleep deprivation was achieved either manually by administering novel objects and occasionally running wheels to the animals or automatically by using a Pinnaclet sleep deprivation apparatus consisting in a moving bar gently sweeping the cage floor, which forced the animal to move and be awake. For the experiments using the novel objects method, both sleeping and sleep deprived mice had access to a running wheel, and one to two unfamiliar objects during their dark period to enhance their environment and facilitate the light/dark entrainment of the rest/activity cycle. During the light period, the group allowed to sleep had their access to running wheels and novel objects removed, whereas these environmental enrichments were retained for the sleep-deprived mice undergoing the sleep deprivation procedure. For the experiments using the automatic apparatus, bar movement speed was set to 4 rpm (four bar sweeps every minute).

#### Sleep and wake assessment via video monitoring

In animals used for molecular and morphological experiments, sleep and wake were monitored with infrared cameras. To estimate sleep and wake, we used a custom-made algorithm to measure motion in these animals. While this method cannot differentiate between NREM and REM sleep time, it reliably distinguishes sleep and wake with a concordance with electroencephalographic (EEG) recordings of about 90% (*27*). Analysis of motion as a proxy for sleep and wake detection prevented the implant of EEG electrodes that inevitability provokes inflammation and can alter subsequent molecular or morphological analysis.

#### Cyclodextrin treatment

2-hydroxypropyl-β-cyclodextrin (CD) was dissolved in sterile saline and administered subcutaneously at a concentration of 2g per kg body weight (*17*). Rats were treated with three injections of CD administered during the dark phase (at 10 am) at day 2, 5, 8 of the 10 days SL experiment.

#### MRI experiments

Rats were divided into a sleep loss (SL, n=10) and sleeping group (S, n=10). The SL group received 10 days of sleep restriction using the moving bar procedure as described before, while the S group experienced the bar movement only for 3 hours/days during the dark period. After the experiment, all rats were perfused transcardially under anesthesia with 0.9% sodium chloride solution followed by 4% paraformaldehyde in PBS. Brains were extracted and post-fixed in the same fixative solution for 2h. After post-fixation, brains were rinsed in PBS and shipped to the MRI facility, where they were included in a falcon with 3% agarose. MRI was performed on a 7 T scanner (Bruker, BioSpect 70/30, Ettlingen, Germany) featuring maximum gradient intensity of 700 mT/m. Diffusion Weighted Magnetic Resonance Imaging (DW-MRI) data were acquired using a stimulated echo planar imaging diffusion sequence, with 126 uniform distributed gradient directions, b=0(6), 4000(60) and 7000(60) s/mm^2^, diffusion time 15 ms, diffusion duration of 5.5 ms, repetition time (TR) = 5000 ms and echo time (TE) = 28 ms. 30 slices were set up to cover the whole brain with field of view (FOV) = 25×25 mm^2^, matrix size = 110 × 110, in-plane resolution = 0.225×0.225 mm^2^ and slice thickness = 0.6 mm. In addition, a T2-weighted image was also acquired with the same resolution to facilitate distortion correction and registration to the rat brain MRI atlas. The total acquisition time per subject was 12 hours.

DW-MRI data were corrected for distortions using linear registration, then processed according to standard diffusion tensor analysis with the software ExploreDTI (*28*), and for CHARMED analysis (*13*) using custom scripts. As such, maps of fractional anisotropy (FA) and restricted signal fraction (RF) were generated for each subject. FA maps were employed to initialize the first steps of an improved version of the TBSS (*29*). This version performs the co-registration steps using Ants normalization package (*30*). We tested for a general linear model comparing SL and control animals, while correcting for multiple comparisons across clusters by using threshold-free cluster enhancement.

#### Histology

After MR analysis, brains were included in 3% agarose/PBS (Sigma‒Aldrich, Madrid, Spain) and cut in a vibratome (VT 1000S, Leica, Wetzlar, Germany) into 50 μm thick serial coronal sections. Coronal sections were rinsed and permeabilized three times in 1x PBS with Triton X-100 at 0.5% (Sigma‒Aldrich, Madrid, Spain) for 10 minutes each and then blocked in the same solution with 4% bovine serum albumin (Sigma‒Aldrich, Madrid, Spain) and 2% goat serum donor herd (Sigma‒Aldrich, Madrid, Spain) for 2 hours at room temperature. The slices were then incubated overnight at 4°C with primary antibodies against myelin basic protein (1:250 Millipore Cat# MAB384-1ML, RRID:AB_240837), and NeuN (1:250, Millipore Cat# MAB377, RRID:AB_2298772) to label myelin and nuclei, respectively. The sections were subsequently incubated in specific secondary antibodies conjugated to the fluorescent probes, each at 1:500 (Molecular Probes Cat# A-11029, RRID:AB_2534088; Molecular Probes Cat# A-11042, RRID:AB_2534099) for 2 h at room temperature. Sections were then treated with 4′,6-Diamidine-2′-phenylindole dihydrochloride at 15 mM (DAPI, Sigma‒Aldrich, Madrid, Spain) for 15 minutes at room temperature. Finally, sections were mounted on slides and covered with an anti-fading medium using a mix solution 1:10 Propyl-gallate: Mowiol (P3130, SIGMA-Aldrich, Madrid, Spain; 475904, MERCK-Millipore, Massachusetts, United States). For myelin labeling, antigen retrieval was performed in 1% citrate buffer (Sigma‒Aldrich, Madrid, Spain) and 0.05% Tween 20 (Sigma‒Aldrich, Madrid, Spain) warmed to 80°C for protein unmasking.

The tissue sections were then examined using a computer-assisted morphometry system consisting of a Leica DM4000 fluorescence microscope equipped with a QICAM Qimaging camera 22577 (Biocompare, San Francisco, USA) and Neurolucida morphometric software (MBF, Biosciences, VT, USA). Myelin fluorescent analysis was performed using Icy software (*31*). Values were then analyzed with a linear mixed effect model using rat as random variable and condition and brain region as fixed variables. The parameters in the LME models were estimated based on maximum likelihood using the lme4 package in R (*32*). Likelihood ratio test was used to assess the statistical significance of the brain region and condition effects.

#### Human data from HCP repository

We used the pre-processed data provided by the Human Connectome Project (www.humanconnectome.org). The used dataset includes subject with DW-MRI scans, the latest pre-processing release (v3.19.0), and a complete behavioral assessment, for a total of 185 subjects. All acquisition parameters and processing pipelines are described in detail on the project page.

We fed the pre-processed images to standard diffusion tensor analysis and to CHARMED analysis pipeline (*13*) using Microstructure Diffusion Toolbox (MDT https://github.com/robbert-harms/MDT). As such, maps of FA and RF were generated for each subject. The same TBSS approach used in animals was used for the analysis, and skeletonized maps were associated voxel-wise with Pittsburgh Quality Index (PSQI) using a general linear model accounting for age, sex, and intra-cranial brain volume, while correcting for multiple comparisons across clusters by using threshold-free cluster enhancement.

#### Electroencephalography

Under deep isoflurane anesthesia (1–1.5% volume), rats were implanted bilaterally for chronic EEG recordings with epidural screw electrodes over the frontal (from bregma: anteroposterior 1 mm, mediolateral, 1 mm) and parietal cortex (anteroposterior, 2 mm; mediolateral, 2 mm) and cerebellum (reference electrode and ground). Electrodes were fixed to the skull with dental cement. Two stainless steel wires (diameter 0.4mm) were inserted into neck muscles to record the EMG. After surgery, rats were housed individually to allow recover. Approximately 7 d were allowed for recovery after surgery, and recordings started only after the sleep–wake cycle had completely normalized. Rats were connected by a flexible cable to a commutator and recorded continuously for 2 weeks using an OpenEphys recording system. EEG and EMG signals were filtered (EEG: high-pass filter at 0.1 Hz; low-pass filter at 40 Hz; EMG: high-pass filter at 10 Hz; low-pass filter at 70 Hz). All signals were sampled at 1000 Hz resolution and down sampled at 512 Hz for analysis. Waking, NREM sleep, and REM sleep were manually scored off-line (SleepSign, Kissei COMTEC, Matsumoto, Japan) in 4-s epochs according to standard criteria. NREM and REM episodes were defined as episodes of duration >= 2 epochs (8-s). One-way ANOVA repeated measures (rANOVA) with factor “day” was performed followed by Dunnett’s multiple comparisons post hoc tests (significance level α=0.05) to evaluate statistical difference against baseline. In case of missing values, data were analyzed by fitting mixed-effects models.

To minimize the number of rats that had to be surgically implanted, these animals were also used to carried out the interhemispheric synchronization analysis. In these analyses, we compared baseline data with those obtained during the second day of recovery. We utilized the second recovery day to let the sleep pressure from CSR subside, which typically eased off during the first recovery sleep day. This approach enabled us to compare days (both baseline and the second recovery day) that exhibited analogous sleep-wake patterns.

##### Interhemispheric synchronization analysis

EEG signals were filtered between 0.15 and 40 Hz by using a FIR (finite impulse response) filter (–6 dB at cutoff frequency, mne.filter function in mne python (*33*, *34*)). Interhemispheric time-domain correlation between EEG amplitude (µV) of the parietal left and right derivation was computed by means of Pearson product-moment correlation coefficient and cross-correlation. Pearson product-moment correlation coefficient for each stage (NREM, REM and WAKE) was calculated to evaluate how the two signals (EEG channels: parietal left, parietal right) co-vary over time, considering the average of all 4-s epochs of NREM, REM and WAKE.

Pearson coefficient was calculated as

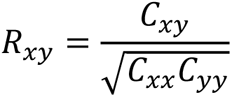

where *C*_*xx*_ and *C*_*yy*_ are the variance and *C*_*xy*_ the covariance matrix of two signals x and y respectively (*35*, *36*).

In addition, we computed mean cross-correlation (*36*) of parietal left and parietal right signals by averaging the values of all 4-s epochs of NREM, REM and WAKE. To analyze the coherence in the frequency domains, we used the coherence function defined as the squared cross-spectrum between signals divided by the product of the auto-spectra of each signal (magnitude squared coherence, MSC). Coherence values ranged from 0 to 1, with 1 indicating full correlation (synchronization) between the two signals at a given frequency, while a value near zero suggesting that the signals were unrelated. The coherence value of two signals x and y, *M*_*xy*_(*f*), was calculated as a function of the spectral densities of signal x, Pxx (f), and y, Pyy (f), and the cross spectral density of x and y, Pxy (f):

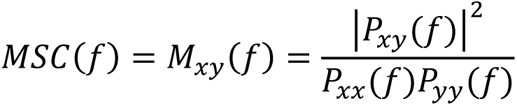

Coherence spectra were calculated between parietal left and parietal right EEG channels for each 4-s epoch considering NREM, REM and WAKE staging. Mean value of coherence was calculated averaging all values of 4-s epochs for each stage respectively. The coherence analysis was performed over specific frequency bands (delta, 0–4 Hz; theta, 4–9 Hz; alpha, 9-12 Hz; sigma, 12– 15 Hz; beta, 15–25 Hz; gamma, 25-50 Hz) by averaging the MSC function in the corresponding frequency range.

Statistical significance was set at *p* < 0.05. To quantify the phase synchronization between two EEG channels, we computed the Phase-Locking Value (PLV). PLV is the most commonly used phase interaction measure, evaluating the absolute value of the mean phase difference between the two signals (*37*). It quantifies the degree to which the phases of two signals are coupled together. PLV ranges from 0 to 1, where 0 indicates no phase locking (completely random phase relationship), and 1 indicates perfect phase locking (both signals have the same phase). PLV between signals *y*_1_(*t*) and *y*_2_(*t*) was computed as follow:

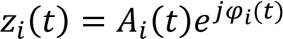

The two analytic signals *z*_*i*_(*t*), were obtained from *y*_*i*_(*t*) (for *i* = {1,2}) using the Hilbert transform (HT):

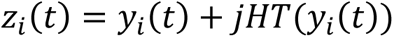

Where *HT*(*y*_*i*_(*t*)) is the Hilbert transform of *y*_*i*_(*t*), which transforms the real signals into a complex representation. Next, we computed the phase difference Δ*φ*_*i*_(*t*) between them (*z*_*i*_(*t*)), with PLV being defined as 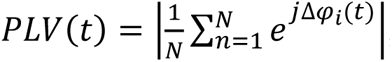, where n indexes the epoch number and N is the total number of epochs (for further details on method see (*37*, *38*)).

In our case, the signals were band-pass filtered to delta-theta frequency range (0.5-9 Hz). Data were expressed as means and standard deviations and tested for normality to ensure they met assumptions of parametric analysis. Paired t-test was performed to evaluate differences in correlation coefficient, MSC and PLV among baseline and second day of recovery in each stage (NREM, REM, WAKE). Statistical significance was set at *p* < 0.05.

#### Cortico-cortical stimulation

Under general anesthesia, rats (n=31) were surgically implanted with LFP electrodes in the primary motor area M1 (AP +2, ML +2) and in the contralateral cortex (AP +2, ML −2). For EEG monitoring, 2 EEG screws were located over the fronto-parietal cortex (AP +4, ML −2 and AP −4, ML −3). The ground and reference were anchored on the cerebellum (Ap - 10, ML +2). EMG electrodes were firmly attached to the superficial muscle at the back of the neck. After 2 weeks of recovery from surgical procedures, animals were divided into sleep loss (SL), sleeping (S), stress, and cyclodextrin (CD) groups. The SL group received 10 days of sleep restriction using the moving bar procedure as described before, while the S group experienced the bar movement only for 3 hours/days during the dark period. In this way, both groups of rats experienced the movement of the bar of the sleep restriction apparatus, thus minimizing possible biases due to the experimental setup. The stress group was subjected to 3hrs of unpredictable chronic mild stress paradigm (UCMS) for 10 consecutive days during the dark phase. UCMS involves exposing animals to systematic and repeated, unpredictable, and uncontrollable stressors over a period of time, leading to behavioral alterations (*39*, *40*). Stressors included exposition to no water, no food, wet bedding, tilting of housing cage (45°), and high frequency rocking (110 RPM for 25sec with an interval of 90sec) for 4 hours at the beginning of the dark cycle. These stressors were repeated twice in succession over 10 days. The CD group received 10 days of sleep restriction using the moving bar procedure as described before and three subcutaneous injections of CD. All animals were allowed to acclimatize to the recording set-up placed in a Faraday box (to limit noise during LFP recording) and to the sleep deprivation chamber for 2 hours before the recording. After acclimatization sessions, three sessions of recordings were performed: 1) a pre-stimulation session to test the input-output signal, signal to noise ratio of the inner/outer channels of the LFP electrodes and to determine optimal stimulation intensity for each animal; 2) baseline before the sleep manipulation experiment (10 days of S, SL, Stress, or SL+CD); 3) Post experiment session after the end of the sleep manipulation experiment.

Electrical stimuli were delivered with an isolated stimulator (ISO-flex; A.M.P.I., Israel) as square-wave pulses (100 ms, 200 μA) and recorded with an OpenEphys acquisition system with a sampling rate of 30KHz. All stimulation occurred in quiet wake. To avoid confounding effects of behavioral state on evoked responses, we carefully monitored the behavior of each rat using LFPs, muscle activity and direct visual observation, and recorded the evoked LFPs under standardized conditions of quiet wakefulness and at the same time of day. All rats were recorded prior (baseline session) and after the sleep manipulation (post session). Notably, post stimulation sessions occurred always 12h after the end of the sleep manipulation to allow the rats to properly recover from the physical activity associated with the sleep restriction procedure. Pre-stimulation session consisted in 20 stimulations/intensities (ranging between 1-60 volts, with a train interval of 15 mins) to carry out a stimulation-response curve used for detecting optimal stimulation intensity. For the baseline and post experiment sessions, 100 pulses (100 ms, 200 μA) at the selected intensity were delivered and the early component of the LFP evoked response was recorded contralaterally. We focused on the transcallosal response because the corpus callosum consists of a distinct, isolated and homogenous bundle of myelinated excitatory fibers, and thus the early monosynaptic component of the evoked response can be easily identified. The response consisted of a depth-negative wave with latency to the peak of ∼ 4-5 ms. Latency was stable within each session and showed minimal (<1ms) variability. The peak latency, amplitude, and slope of the evoked LFP response were estimated in custom-made MATLAB routine. The slope of the component was computed as mean first derivative of the first down-going segment. Repeated measures ANOVA with time as within factor and group as a between factor was used to evaluate differences between baseline and post experiment sessions. Šídák’s multiple comparisons test was used as a pot-hoc test. After the last stimulation session, rats were perfused transcardially under anesthesia with 0.9% sodium chloride solution followed by 4% paraformaldehyde in PBS and the brain was harvested for histology. Histological evaluation was performed to validate the position of the LFP electrodes. After harvesting the perfused brain, the brain was post-fixed in 4%PFA for 72 hours and 50-μm serial coronal section of the brain slice was collected with vibratome. Sections were stained with Nissl dye and the position of the LFP electrode of all experimental animals was located within the primary motor cortex M1.

#### Gene expression analysis

we used the array data available at NCBI GEO database ((*6*); GSE48369) to perform gene expression analysis of forebrain samples collected from sleeping and sleep deprived mice. Samples of this database were collected using the genetically targeted translating ribosome affinity purification (TRAP) methodology from BAC transgenic mice expressing EGFP tagged ribosomal protein L10a in oligodendrocytes. This method permits to study the expression of mRNAs attached to ribosomes on their way to become proteins, thus providing a better functional overview than traditional methods based on estimates of whole RNA. In the present study, we used array data obtained only from samples of mice after undisturbed sleep (˃45 min, discontinued by period of wake of ˂4 min, S) and after a period of 4 hours of sleep loss (SL) obtained with the exposure to novel objects. Data were normalized within each behavioral state group using Robust Multiarray Average. To identify transcripts that were differentially expressed across S and SL, comparisons were carried out using the Welch’s t test with Benjamini and Hochberg false discovery rate (FDR) multiple-test correction. The lists of differentially expressed genes was submitted to the DAVID (Database for Annotation, Visualization and Integrated Discovery) bioinformatics database for functional annotation (http://david.abcc.ncifcrf.gov/) (*41*). The background list used in the program included all the genes assigned to Affymetrix GeneChip Mouse Genome 430 2.0 arrays.

#### Myelin tissue collection and preparation

Sleeping mice (S) were euthanized during the light phase (at ∼3.00-5.00 PM) following a long period of sleep (˃45 min, discontinued by period of wake of ˂4 min) and after spending no less than 70% of the previous 6–7 h awake. In the sleep loss (SL) group, 6-8 hours of sleep deprivation was manually performed by an experimenter during the light phase by exposing the mice to novel objects and occasionally to a running wheel when animals appeared drowsy. Mice were never disturbed during eating or drinking. Mice were sacrificed with cervical dislocation, their brain quicky removed and snap frozen. To obtain myelin preparations we followed a modified version of the LaRocca’s protocol (*42*). Briefly, brain tissue was weighted and then homogenized on ice using a glass potter containing in 0.3 M sucrose solution with 20mMTris·Cl buffer (pH 7.45), 1mM EDTA, 1mM DTT, 100 μM phenylmethylsulfonylfluoride (PMSF), 10 μg/ml leupeptin, and a mixture of anti-proteolytic compounds (cOmplete Tablets, Roche). The homogenate was layered over 0.83 M sucrose solution and centrifuged 35 min at 75,000×g. The band of crude myelin membranes formed at the 0.3 M /0.83 M sucrose interface was collected, and after washing out the sucrose with hypotonic buffer, was subjected two times to a cycle of hypoosmotic shock and low-speed centrifugation (15 min at 12,000 × g) to remove cytoplasmic and microsomal contaminants. Then myelin was further purified with a repetition of the first density gradient centrifugation and a cycle of hypoosmotic shock and low-speed centrifugation. Finally, a highly-purified myelin fraction was prepared by a third density gradient centrifugation. However, in this step myelin was resuspended in 0.83 M sucrose, and the 0.83 M sucrose solution was laid over with 0.30 M sucrose. The myelin fraction at the interface was subjected to a final hypoosmotic shock cycle, collected, and resuspend in 500 µl of Tris·Cl buffer. Protein concentration was assessed using a spectrophotometer.

#### LC-MS/MS lipidomic analysis

Highly-purified myelin fractions (200µl) from S (n=10) and SL (n=8) animals were extracted with MTBE/MeOH/H2O (10:3:2.5) according to Matyash et al (*43*). Briefly, to each sample (in a 2 ml tube) were added MeOH (290 µl) and the IS pool (10 µl MeOH). After vortexing 1 ml of MTBE was added, and the sample sonicated and vortexed again, was allowed to extract at 10°C under shaking. After 1 hr, 250 µl of milliQ water were added to induce phase separation. After 10 min shaking, the sample was centrifuged at 10,000 g for 10 min at 4°C and the upper phase was recovered. The extraction was repeated by adding 300 µl MTBE and the organic extract recombined, dried under nitrogen stream and vacuum, and stored at −80°C until analyses. For LCMS, samples were reconstituted in 200 µl MeOH/isopropanol (1:1).

Chromatographic separations were achieved on Infinity 1290 UHPLC System (Agilent Technologies, Santa Clara, CA, USA), equipped with a Kinetex Biphenyl 2.6 µm, 150 x 2.1 mm column, (Phenomenex, Castel Maggiore, Bologna, Italy) according to Cutignano et al. (*44*) with minor modifications. Briefly, eluent A was acetonitrile/H_2_O 60:40, 10 mM ammonium formate, 0.1% FA and eluent B: isopropanol/acetonitrile 90:10, 2 mM ammonium formate, 0.1% FA. All solvents were LC-MS grade. The elution program consisted of a gradient from 20 to 40% B in 6.5 min, then to 50% B up to 13 min, reaching 90% B at min 16, holding for 1 min and returning back to 20% B in 1 min. A post run equilibration step of 5 min was included prior to each analysis. Column temperature was set at 40°C. Flow rate was 0.3 ml/min. The injection volume was 5 μl and the autosampler was maintained at 10°C.

MS analyses were carried out on Q-Exactive Hybrid Quadrupole-Orbitrap mass spectrometer (Thermo Scientific, San Jose, CA, USA) equipped with a HESI source. Source parameters were as follows: spray voltage positive polarity 3.2 kV, negative polarity 3.0 kV, Capillary temperature 320°C, S-lens RF level 55, Auxiliary gas temperature 350°C, Sheath gas flow rate 60, Auxiliary gas flow Rate 35. Full MS scans were acquired in the range 150–1800 m/z at 70000 of mass resolution, AGC Target 1e6, Acquisition time 100ms. For MS/MS analysis a data dependent ddMS2 Top10 method was used; Mass Resolution was 17500, AGC Target 1e5, Acquisition Time 75 ms. Mass fragmentation was obtained with a stepped normalized energy (NCE) of 16–20-30 and 20–40 in positive and negative ionization mode, respectively. A pool of commercial and in-house synthesized standards was used as Internal Standard mix for quantitative purposes. After preliminary experiments run to assess the appropriate concentration, the customized mix included the following (final concentration in parentheses): Triacylglycerol 17:0/17:0/17:0 (1 µg/ml), Diacylglycerol 15:0/18:1 (0.1 µg/ml), Phosphatidyl choline 10:0/10:0 (2 µg/ml), Phosphatidyl choline 15:0/18:1 (2 µg/ml), Phosphatidyl ethanolamine 17:0/17:0 (2 µg/ml), Phosphatidyl serine 17:0/17:0 (2 µg/ml), Phosphatidyl inositol 15:0/18:1 (2 µg/ml), Phosphatidyl glycerol 17:0/17:0 (2 µg/ml), Lysophosphatidyl choline 17:0 (0.2 µg/ml), Lysophosphatidyl ethanolamine 17:1 (0.2 µg/ml), Ceramide d18:1/17:0 (0.5 µg/ml), Glucosyl ceramide d18:1/17:0 (0.2 µg/ml), Plasmalogen 18:1d9 (2 µg/ml), Sphingomyelin d18:1/17:0 (1 µg/ml), Sulfogalactosyl ceramide d18:1/17:0 (0.5 µg/ml), MGDG 19:0/19:0 (1 µg/ml). All standards were purchased from Avanti Polar Lipids. except MGDG which was synthetized in house as reported in (*45*).

Each analysis was run in triplicate. Raw LC-MS/MS data were processed by Xcalibur software (Thermo Scientific, version 3.1.66.10); lipid species were identified with the support of LipidSearch software (Thermo Scientific, version 4.1.30). A tolerance of 5 ppm was set for Precursor Ion and 10 ppm for Product Ion. The m-score threshold was set to 5. The lipid identification lists were aligned for control and treated samples and compared by their lipid class and lipid species levels using a retention time tolerance of +/- 0.25 min. The main grade was set to A, B and C for all lipid classes. All data were manually double checked. Data were analyzed with LipidSig to assess the quality and the clustering of samples and quantify differentially expressed lipids (*46*). Absolute quantitative data were reported as µg lipid/ml myelin sample. Differentially expressed analysis was performed to find significant lipid species. Sample expression data were analyzed with unpaired t-test with p-value adjusted by Benjamini-Hochberg procedure.

#### LC-MS analysis of cholesterol

Highly-purified myelin fractions (200µl) from S (n=10) and SL (n=8) animals were extracted with a modified Bligh and Dyer protocol (*47*). Briefly, samples were sonicated and then extracted with chloroform/methanol (2:1, v/v) containing internal deuterated standard for cholesterol quantification by isotope dilution (2 ug/ml for d7 cholesterol, Merck). Organic phases were collected and dried down under nitrogen. Then lipid extracts were analyzed by liquid chromatography-atmospheric pressure chemical ionization-single quadrupole mass spectrometry (LC-MS2020 Shimadzu). Briefly, using APCI positive ionization, cholesterol was acquired in SIM mode with a m/z of 369.5 and 376.5. MS parameters were the following: acquisition time 0-15, event time 0.5 sec, detector voltage 1.7 kV, interface temperature 400 °C, DL temperature 250 °C, heat block 230 °C, nebulizing gas flow 3 L/min and drying gas 5 L/min. A Phenomenex Kinetex C18 (5 μm x 4.6 mm x 150 mm) column was used for isocratic elution utilizing 95% Mobile Phase B for 15 minutes at a flow rate of 0.5 mL/min. Cholesterol eluted off the column with a retention time of 5.54 min. The column temperature was 40°C and an injection volume of 10 μL was used. Mobile Phase A consisted of 0.1% formic acid in DiH2O while Mobile Phase B contained 0.1% formic acid in acetonitrile. Endogenous levels of cholesterol were calculated based on their area ratio with the internal deuterated standard signal areas and normalized to ml of myelin. Differences between groups were analyzed with unpaired t-test.

#### Membrane fluidity

Physico-chemical studies of highly-purified myelin fractions from S (n=6) and SL (n=6) mice were performed by using two fluorophores: 2 dimethylamino lauroyl naphtalene (Laurdan), located at hydrophobic hydrophilic interface of the membrane, and 1,6 diphenyl 1,3,5 hexatriene (DPH), incorporated at different levels of the membrane hydrophobic core. The quantum yields of fluorescence of both probes are virtually zero in aqueous solutions, while are quite high in membranes.

DPH fluorescent anisotropy is widely used to study the organization and dynamics of the internal regions in membranes (*48*). It gives information about membrane fluidity that is a measure of the bilayer resistance to rotational and translational motions of molecules and reflects lipid packing in the bilayer.

The fluorescence anisotropy was calculated by using the following equation:

rs = (I‖ - IꞱxg)/(I‖ + 2IꞱxg) where g is an instrumental correction factor, I‖ and IꞱ are, respectively, the emission intensities with the polarizers parallel and perpendicular to the direction of the polarized exciting light.

Laurdan fluorescence excitation and emission spectra are affected by the amount of water (polarity) and by the motion of water molecules (dipolar relaxation of water molecules) close to the fluorescent moiety. Spectroscopic data were used to calculate the excitation and emission generalized polarization spectra, which provide information about the lipid packing (fluidity) of the membrane and the phospholipid phase (*49*). Laurdan excitation GP (Ex GP) and emission (Em GP) spectra have been calculated as follow (*50*):

Ex GP=(I440-I490)/(I440+I490) where I440 and I490 are the intensities at each excitation wavelength, from 320 to 420 nm, obtained using a fixed emission wavelength of 440 and 490 nm, respectively; Em GP=(I390–I360)/(I390+I360) where I390 and I360 are the intensities at each emission wavelength, from 420 to 550 nm, obtained using a fixed excitation wavelength of 390 and 360 nm, respectively. Stock solutions of Laurdan (6-dodecanoyl-2-dimethylamine-naphthalene) in ethanol and DPH (1,6-diphenyl-1,3,5-hexatriene) in tetrahydrofuran were added to myelin membranes at final probes concentration of 1 μM; each suspension was incubated in the dark, at 37 ◦C for 2 h prior to use. The fluorescence measurements have been performed at 37°C with a computer-controlled PerkinElmer LS55 spectrofluorimeter. The temperature was measured in the sample by a digital thermometer. The background fluorescence of the samples was checked prior to each measurement and was negligible when the probes were added. Differences between groups were analyzed with unpaired t-test for DPH and 2-way ANOVA with wavelength and conditions as factors for Laurdan measurements.

#### BODIPY-cholesterol assessment

Rats were divided into a sleep loss (SL, n=5) and sleeping groups (S, n=5). The SL group received 10 days of sleep restriction using the moving bar procedure as described before, while the S group experienced the bar movement only for 3 hours/days during the dark period. In another set of experiments, we compared SL treated with cyclodextrin (SL+CD, n=5) and sleeping (S, n=5) rats. After the experiment, all rats were perfused transcardially under anesthesia with 0.9% sodium chloride solution followed by 4% paraformaldehyde in PBS. Brains were post-fixed for 48 h and then sliced into 40 µm sections using a vibratome.

After blocking with normal goat serum (NGS) 10% with 0.05%TritionX-100, frontal sections were incubated with Bodipy-cholesterol (TopFluor, Avanti Polar Lipids) at 8 µg/ml in association with myelin basic protein antibodies (MBP, 1:1000, Abcam Cat# ab65988 RRID:AB_1139419) at room temperature (2h) and then at 4°C overnight. After washing with PBS, sections were incubated with secondary antibodies (1:1000) and mounted onto slides for microscopy analysis. Confocal fields (3 per section, 2 sections per rat) were acquired in the corpus callosum. MBP staining was used to narrow the analysis on myelin enriched regions. Images contained mostly myelinated fibres and sporadic cells that were outlined and not included in the analysis. To estimate cholesterol levels, mean 488 nm fluorescence intensity was calculated for each image using FiJI. To estimate BODIPY-cholesterol levels in neurons and astrocytes, S and SL frontal sections were first incubated with a blocking buffer (bovine serum albumin 3% with 0.3%TritionX-100 for NEUronal Nuclei [NeuN], NGS 10% with 0.05%TritionX-100 for Glial fibrillary acidic protein [GFAP]) for 1 h and then with BODIPY-cholesterol (TopFluor, Avanti Polar Lipids) at 8 µg/ml in association with antibodies against NeuN (1:200, Synaptic system Cat# 266011, RRID:AB_2713971) or GFAP (1:500, Sigma, Cat# G3893, RRID:AB_477010) overnight at 4°C. After washing with PBS, sections were incubated with secondary antibodies (1:1000) and mounted onto slides for microscopy analysis. Confocal fields (3 per section, 2 sections per rat) were acquired in the lower layers of the frontal cortex. To estimate cholesterol levels in neurons and astrocytes, somas of NeuN and GFAP positive cells were manually segmented and mean 488 nm fluorescence intensity referring to BODIPY-cholesterol was measured within each individual cell using FiJI.

For all these experiments, values were analyzed with a linear mixed effect model using rat as random variable and condition as a fixed variable. The parameters in the LME models were estimated based on maximum likelihood using the lme4 package in R (*32*). Likelihood ratio test was used to assess the statistical significance of the condition effect.

#### Rotarod experiment

Rats were divided into a sleep loss (SL, n=12), a sleeping (S, n=12), a SL group treated with Cyclodextrin (SL+CD, n=9). The SL group received 10 days of sleep restriction using the moving bar procedure as described before, while the S group experienced the bar movement only for 3 hours/days during the dark period. The CD group received 10 days of sleep restriction using the moving bar procedure as described before and three subcutaneous injections of 2-hydroxypropyl-β-cyclodextrin 2 g per kg body weight dissolved in sterile saline. Notably, to avoid the confounding effect of physical fatigue on the motor performance evaluation, SL rats were given 12 hours of rest during the light cycle before testing: during this period the animals were allowed to sleep and rest ad libitum. To reduce the procedural stress, all rats were manually handled by the same operator that performed the behavioral test and briefly (5 min) exposed to the rotarod apparatus in off-mode for at least two weeks prior to the test. We used an accelerating rotarod system from Ugo Basile SRL. Rats were placed onto a stationary rod and acceleration began. The acceleration profile was linear (starting speed 10 rpm; final speed: 80 rpm; time to accelerate from 10 to 80 rpm: 80 sec). Time when rats fell off the rod were automatically recorded. Sessions included 10 trials for each rat. A motor performance score was calculated for each rat by averaging the scores of each trial, while a motor learning score was computed by dividing the averaged time-to-fall of the last two trials by the time-to-fall of the first trial. Data were analyzed using one-way ANOVA followed by Tukey’s multiple comparisons test.

#### Statistics

statistical methods used to derive significances are described in each section of the previous paragraphs. Computational and statistical analyses were performed by using Python 3.9, R, and GraphPad Prism 9.5.1.

## Supplementary Text

### Correlation of white-matter (WM) integrity and Pittsburgh sleep quality index (PSQI) score

The correlation between diffusion magnetic resonance imaging parameters and PSQI scores was examined in the WM of 185 subjects (from the S1200 Human connectome project (HCP) data release). Age, sex, and intra-cranial brain volume were included as covariates for the analysis. Tract-based spatial statistics (TBSS) revealed a significant negative correlation between fractional anisotropy (FA) derived from diffusion tensor imaging (DTI) of WM regions of the right hemisphere and PSQI scores. Restriction fraction (FR), derived from the CHARMED model, although not significant, tended to be negatively correlated with PSQI score in regions that appear to correspond to those where significant FA-PSQI negative correlations were found (Figure S1). Based on the PSQI scores, subjects were divided into the good sleep (PSQI < 3, N=30) and poor sleep (PSQI > 5, N=51) groups, respectively. No statistically significant differences were observed for age (P=0.48), neither for sex (P=0.29) between the groups. Voxel-wise statistics showed a significant reduction of FA in the poor sleep group (Figure S2A). Accordingly, RF also tended to be reduced in the poor sleep group. While the TBSS-threshold-free cluster enhancement method did not detect significant changes in RF between the groups, the poor sleep group exhibited a decrease in the mean of FR when calculated from regions showing a significant reduction in FA (Figure S2B).

### Automatic sleep restriction leads to effective reduction of sleep time

To test the efficacy of the automatic sleep restriction system, we recorded 6 rats for 7 days with EEG electrodes. During this period, we recorded 24 hours of undisturbed baseline, 4 consecutive days of sleep restriction consisting of 4 hours of sleep opportunity (4S+) in which the apparatus was in OFF mode and 20 hours of sleep deprivation (20S-) during which the sweeping bar was on each day, and 2 days of undisturbed recovery (Figure S3A; representative EEG recordings in Figure S3B).

EEG data analysis showed that the automatic sleep restriction procedure significantly reduced the time spent in NREM and REM sleep with respect to baseline (rANOVA, for NREM F_day_ (2.685, 13.43)=116, P<0.0001, for REM F_day_ (2.438, 12.19)=222.5, P<0.0001), while it increased the time of wake (F_day_ (2.358, 11.79)=188.6, p<0.0001). In addition, it led to rebound sleep, particularly by increasing the duration of REM sleep during the first day of recovery (P=0.0006, Figure S3C).

To gain further insights into the homeostatic response to this sleep restriction approach, the experimental design was set to allow the rats to sleep ad libitum for 4 hours (4S+) each day from SL1 to SL4, while they were sleep restricted by rotating bar during the remaining 20 hours (20S-). During 4S+ the time spent in NREM sleep did not change in SL1 and 2, while it slightly decreased in SL3 and SL4 (rANOVA, for NREM F_day_ (2.097,10.48)=7.040, P=0.0019 relative to baseline, Supplementary Figure 3D). On a fine-grained analysis this effect was associated with reduced number but larger mean duration of NREM episodes in SL2, SL3, SL4 (number: rANOVA, F(4, 20)=47.79, P<0.0001); duration: rANOVA, (4, 20)=11.46, P<0.0001, Supplementary Figure 3E). While the total time spent in NREM sleep was not greatly changed, the time spent in REM sleep remarkably increased in SL2, SL3, and SL4 relative to baseline (F_day_ (2.212,11.06)=34.48, p<0.0001, Figure S3D). During the 20S- the time spent in NREM and REM sleep was strongly reduced, while the time spent in wake was increased for SL1-4 (rANOVA, for NREM F_day_ (2.303, 11.52)=70.70, P<0.0001, for REM F_day_ (1.171,5.857) =137.7, p<0.0001, for wake F_day_ (2.491,12.46)=111.7, P<0.0001, Figure S3F). Of note, the sleep restriction procedure increased the number of NREM attempts in SL1, SL2, SL3 (rANOVA, F (4, 20)=7.233, P=0.0009 relative to baseline) while their related mean duration was largely shorter than the one observed during baseline for all 4 days of sleep restriction (rANOVA, F (4, 20)=111.0, P<0.0001, Figure S3G).

Thus, the automatic sleep restriction system was effective in reducing the overall sleep time by leading to manifest signs of increased sleep pressure.

---

**Fig. S1:**
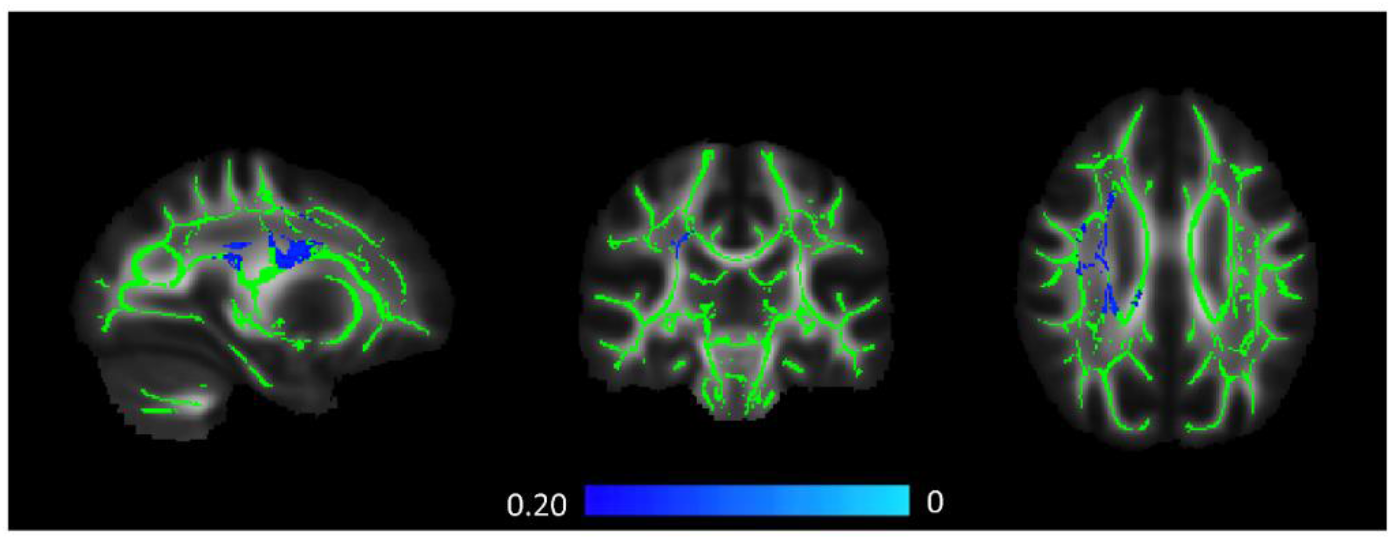
correlation between restricted fraction (RF) and Pittsburgh sleep quality index (PSQI) score. Blue/ light blue represent the white-matter regions, from tract-based spatial statistics (TBSS) with negative correlation between RF and PSQI (n=185). Correlational analyses were conducted using randomise threshold-free cluster enhancement (TFCE), and family-wise corrected for multiple comparisons. The results are shown in overlay on the mean FA template and the mean FA skeleton (green), calculated from all the subjects. Axial and coronal sections are shown with the right hemisphere on the left; sagittal section is shown with posterior on the left.

**Fig. S2:**
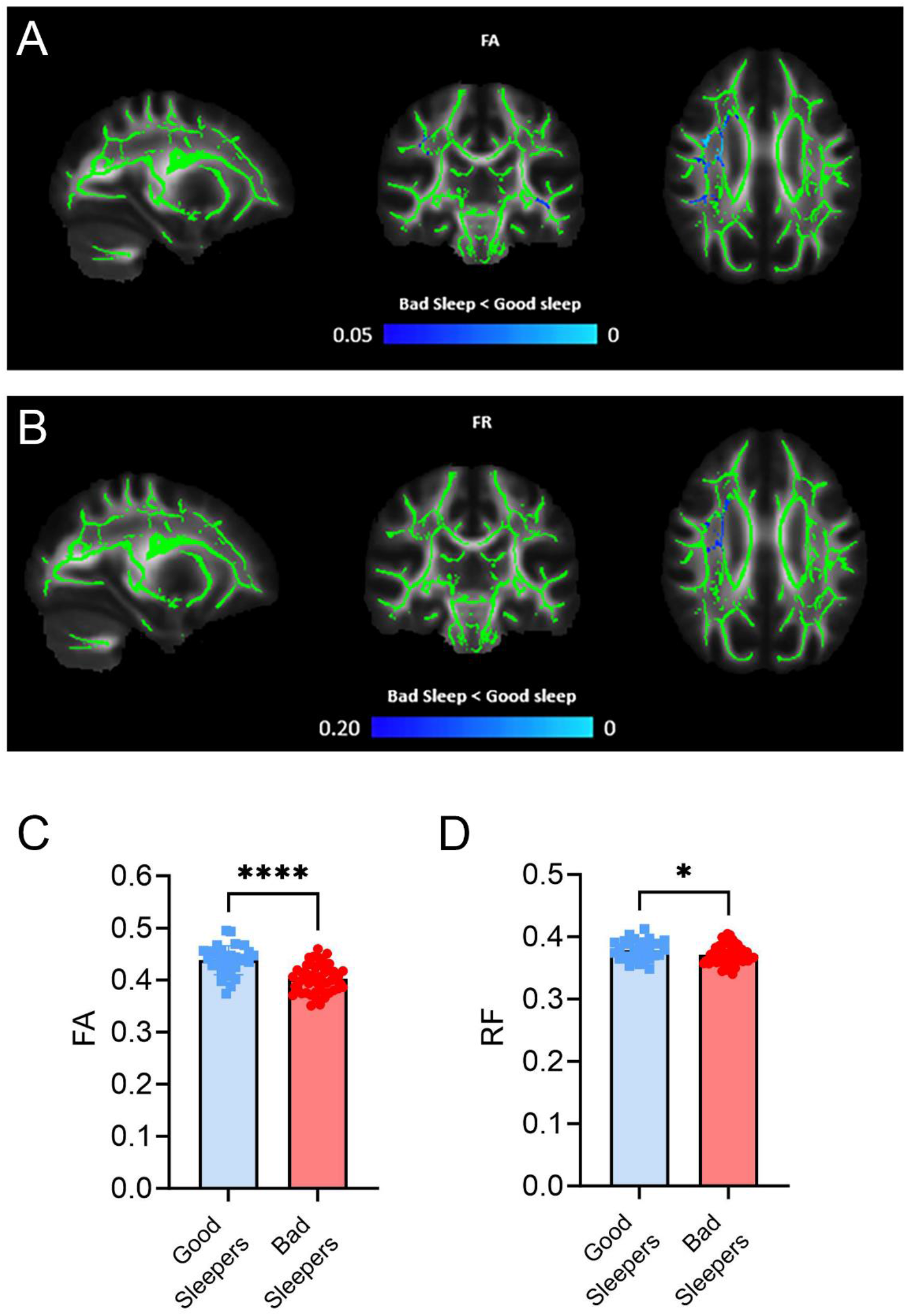
comparison of WM integrity between poor and good sleepers. **A-B.** Fractional anisotropy (FA, **A**) and restricted fraction (RF, **B**) differences between poor (n=51) and good (n=30) sleepers. Blue/ light blue represent the white-matter regions, from tract-based spatial statistics (TBSS), with significantly reduced FA (**A**) and a trend toward the reduction for RF (**B**) in the poor sleep group. Correlational analyses were conducted using randomize threshold-free cluster enhancement (TFCE), and family-wise corrected for multiple comparisons. The results are shown in overlay on the mean FA template and the mean FA skeleton (green), calculated from all the subjects. Axial and coronal sections are shown with the right hemisphere on the left; sagittal section is shown with posterior on the left. **C**. Mean fractional anisotropy (FA) between groups in regions where anisotropy fraction (FA) was significant in TBSS analysis. Statistical significance was addressed with unpaired t-test. ****P<0.0001. **D**. 0001Mean restricted fraction (RF) between groups in regions where anisotropy fraction (FA) was significant in TBSS analysis. Statistical significance was addressed with unpaired t-test. *P<0.05.

**Fig. S3:**
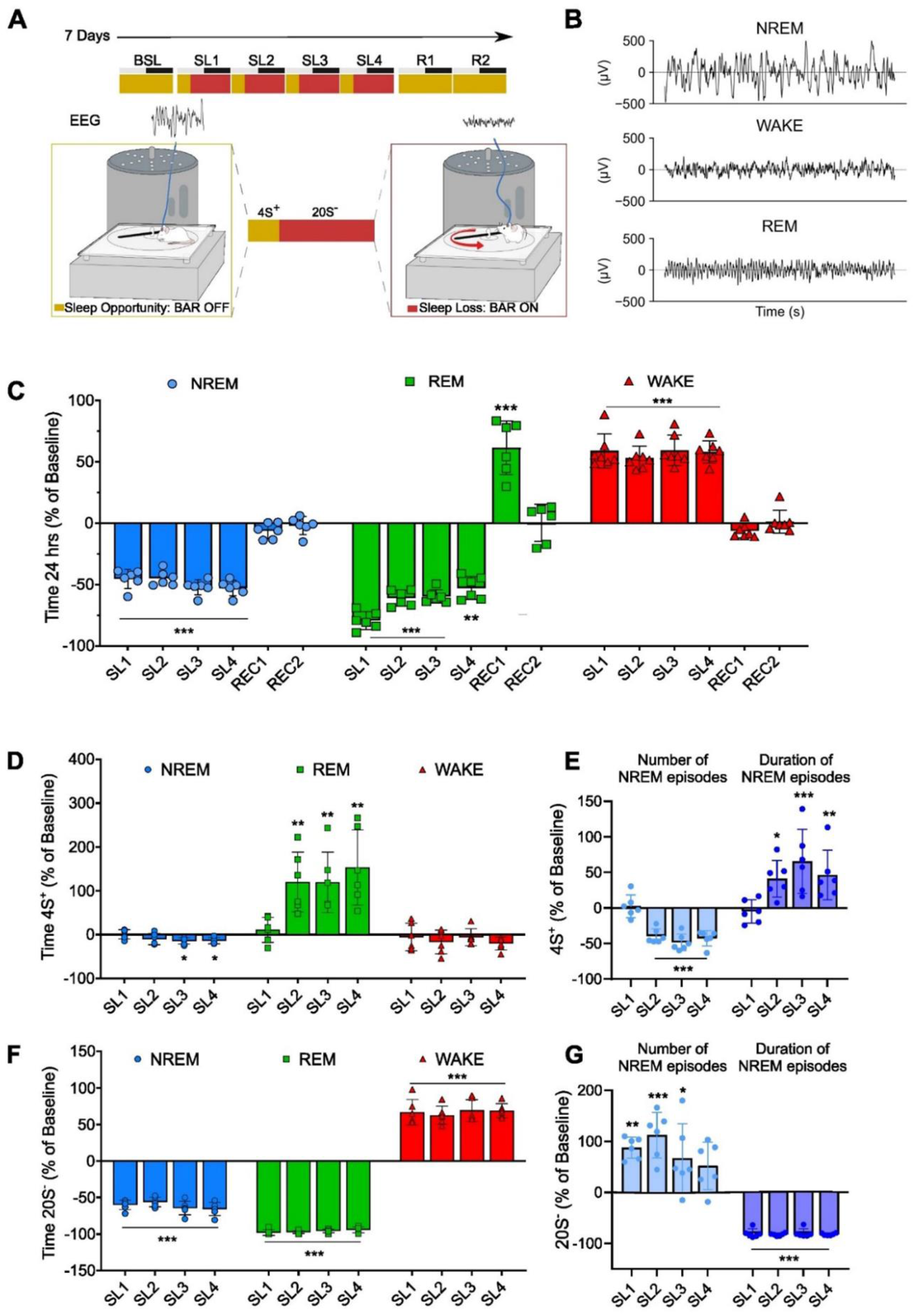
effects of the automated sleep restriction. **A.** Experimental design of EEG experiments (n=6). BSL=baseline, SL= sleep loss, R=recovery from SL. 4S+ (in yellow) indicates 4-hour sleep opportunity, 20S- (in red) indicates sleep restriction. **B.** Examples of EEG traces in WAKE and NREM and REM sleep **C**. Time spent in NREM (blue) and REM (green) and wakefulness (red) over the 24 hours during sleep restriction (SL1-SL4) and post restriction recovery days (REC1-2). **P<0.01; ***P<0.001. **D**. Time spent in wakefulness (red), NREM (blue) and REM (green) sleep during the sleep opportunity window of 4 hrs (4S^+^). **E**. Number and duration of NREM episodes during 4S^+^. **F** Time spent in wakefulness (red), NREM (blue) and REM (green) sleep during the 20 hrs of sleep restriction (20S^-^). **G**. Number and duration of NREM episodes during 20S^-^. For **C-G** values are expressed as % of the baseline. *P<0.05. **P<0.01. ***P<0.001.

**Fig. S4:**
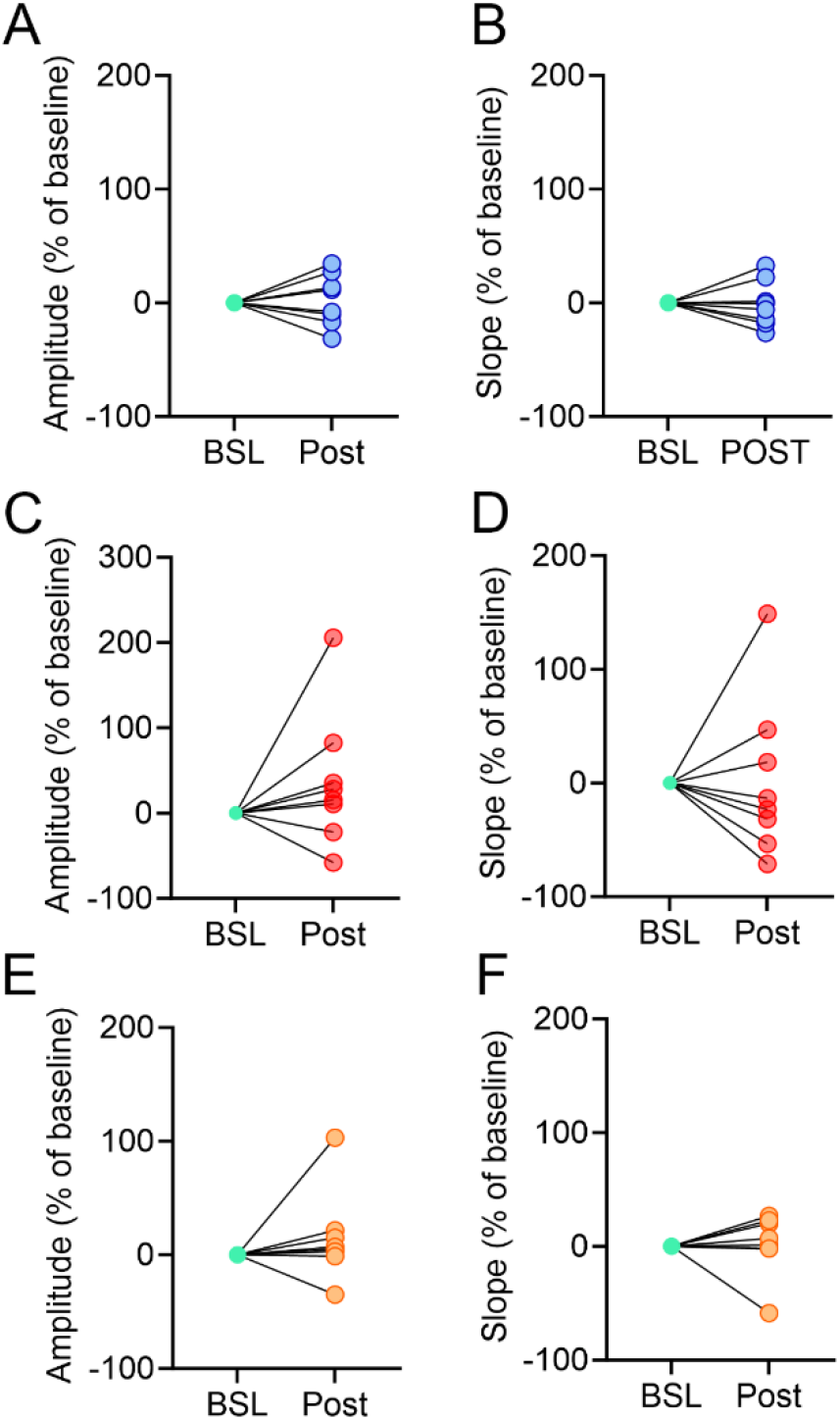
effects of S, SL, and Stress on amplitude and slope of the cortico-cortical transcallosal evoked responses. **A-B.** Amplitude (**A**) and slope (**B**) of the early component negative peak for SL individual rats. **C-D.** Amplitude (**C**) and slope (**D**) of the early component negative peak for S individual rats. **E-F.** Amplitude (**E**) and slope (**F**) of the early component negative peak for Stress individual rats.

**Fig. S5:**
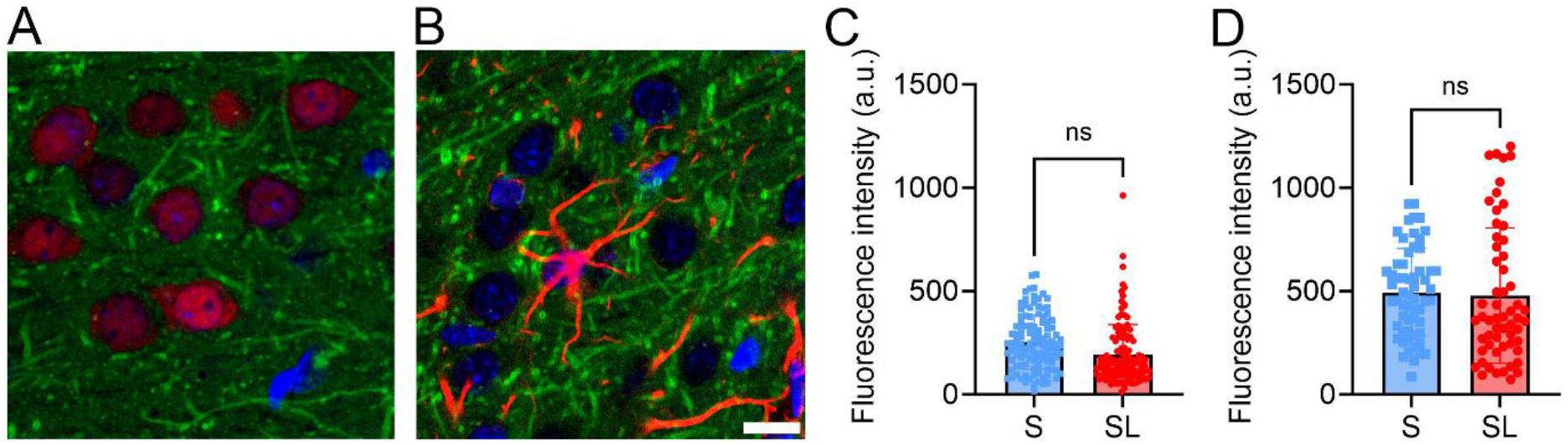
effects of S and SL on BODIPY-cholesterol levels in neurons and astrocytes. **A-B.** Representative images of neurons (Neun+ cells, red in A) and astrocytes (GFAP+ cells in B, red) in association with BODIPY-cholesterol (green) and DAPI (blue). Scale bar: 10 µm. **C-D.** BODIPY-cholesterol levels estimated within the neuronal (**C**) and astrocytic (**D**) soma in S (n=5) and SL (n=5) rats.

